# Methylation content sensitive enzyme ddRAD (MCSeEd): a reference-free, whole genome profiling system to address cytosine/ adenine methylation changes

**DOI:** 10.1101/616532

**Authors:** Gianpiero Marconi, Stefano Capomaccio, Cinzia Comino, Alberto Acquadro, Ezio Portis, Andrea Porceddu, Emidio Albertini

## Abstract

Methods for investigating DNA methylation nowadays either require a reference genome and high coverage, or investigate only CG methylation. Moreover, no large-scale analysis can be performed for N^6^-methyladenosine (6mA). Here we describe the methylation content sensitive enzyme double-digest restriction-site-associated DNA (ddRAD) technique (MCSeEd), a reduced-representation, reference-free, cost-effective approach for characterizing whole genome methylation patterns across different methylation contexts (e.g., CG, CHG, CHH, 6mA). MCSeEd can also detect genetic variations among hundreds of samples. MCSeEd is based on parallel restrictions carried out by combinations of methylation insensitive and sensitive endonucleases, followed by next-generation sequencing. Moreover, we present a robust bioinformatic pipeline (available at https://bitbucket.org/capemaster/mcseed/src/master/) for differential methylation analysis combined with single nucleotide polymorphism calling without or with a reference genome.

## Introduction

DNA methylation is one of the fastest mechanisms that organisms use to rapidly adapt to new conditions^1–3^. Indeed, the methylation of cytosine residues in genomic DNA has a pivotal role in regulation of genome expression^4–7^, particularly for the cytosines in promoter sequences of specific genes. Generally, methylation is correlated with silencing of genes and transposable elements, while demethylation is correlated with active transcription^4^, although the reverse has also been documented^8^. Moreover, methylation patterns along a gene can have specific effects on the gene expression: body-methylated genes tend to be constitutively expressed, whereas promoter-methylated genes are preferentially expressed in a tissue-specific manner^5^.

Cytosine methylation is conventionally classified in terms of CG, CHG, and CHH sequence contexts (where H is A, C, or T), which are subjected to the actions of different DNA methyltransferases^9–12^. Methylation on adenine (N^6^-methyladenosine; 6mA)^7,13^ has been recently found in *Chlamydomonas* and in several multicellular eukaryotes, including flowering plants^7,14–18^. While in prokaryotes and ancient eukaryotes, 6mA serves as major marker to discriminate invasive foreign DNA^19^, its role in eukaryotes has recently been associated with transcriptional activation in response to stress^7,14^, and with transgenerational chromatin regulation^20^.

There are many technologies available to obtain genome-wide information on differential DNA methylation. Some of these provide qualitative information on the methylation state, while others are based on the chemical conversion of unmethylated cytosine to thymine. This chemical conversion defines the level of unconverted cytosines, which provides a measure of the level of DNA methylation^21–23^.

Whole-genome bisulfite sequencing (WGBS) is a technique that can assess virtually every cytosine methylation state in the genome, although this requires high coverage (at least 5-10–fold)^21,24,25^, which makes it expensive in species with a large genome and/or in experiments with many samples^26^. Reduced representative bisulfite sequencing (RRBS)^27^ has been proposed as an alternative to WGBS for large genomes. RRBS introduces a DNA digestion step with a methylation-insensitive enzyme that is followed by size filtration of the restricted fragments, and chemical conversion. This technique investigates only a fraction of the genome, but the increased sequencing coverage of the represented fraction provides greater confidence in such methylation measurements. Although RRBS has been shown to be extremely powerful, there remain technical and financial bottlenecks that challenge the feasibility of these approaches, especially in species with a large genome and lacking a reference genome^27,28^.

Further reductions in sequencing efforts can be achieved through adoption of methylation-sensitive endonucleases, as in methylation-sensitive restriction-enzyme digestion and sequencing (MRE-seq)^29^, or the EpiRADseq^30^ variation of Double digest RADseq (ddRADseq^31^). These techniques involve DNA digestion with a methylation-sensitive enzyme followed by size selection and sequencing. However, the kinetics of these methylation-sensitive enzymes have a bias toward demethylated sites, and thus they act more specifically on hypomethylated sites, which are the most difficult to detect with conventional techniques^29^. Such read counts for each locus do not provide absolute measures of cytosine methylation, although they can be useful to infer methylation differences between samples at specific sites. A limitation of MRE-seq is that it is specific for CG methylation. Importantly, none of these methods address analysis of the methylation status of adenines, which is currently being studied using very costly approaches, such as mass spectrometry, immunoprecipitation, and PacBio sequencing^7,32^.

To overcome some of these limitations, we present the methylation content sensitive enzyme double-digest restriction-site-associated DNA (ddRAD) technique (MCSeEd), a very simple, highly scalable, cost-effective extension of the original ddRAD protocol that allows the detection of methylation changes for the CG, CHG, CHH, and 6mA contexts.

This MCSeEd technique was tested in two maize experimental systems: (i) leaves of a commercial maize hybrid grown under normal irrigation (well watered; WW) and under drought stress (DS), and collected 60 days after sowing (DAS); and (ii) shoots and roots of the inbred line B73 collected at 5 DAS. The relative methylation changes estimated by MCSeEd for differentially methylated positions (DMPs) and differentially methylated regions (DMRs) clearly discriminated between these samples (i.e., WW *vs* DS; B73 shoots *vs* roots) with both genome-dependent and genome-independent approaches. The DMRs identified by MCSeEd showed gene enrichments that were related to the experimental system under investigation. Shifts in single-nucleotide polymorphism (SNP) allele frequencies were also identified, and were related to the specific methylated/ unmethylated alleles.

## Results

### MCSeEd efficiently identifies methylation variations induced by drought stress in maize leaves

#### Differentially methylated positions

The MCSeEd technique was used to monitor DNA methylation changes induced by drought stress in maize leaves. To this end, we constructed next-generation sequencing (NGS) libraries from genomic DNA purified from the leaves of the WW and DS maize plants. A total of 24 libraries were produced by double restriction–ligations, each using *Mse*I in combination with one of the four methylation-sensitive enzymes *Aci*I, *Pst*I, *Eco*T22I, and *Dpn*II, for the CG, CHG, and CHH and 6mA contexts, respectively^29,33–36^ (Supplementary Table S1) as outlined in Figure 1 and in Supplementary Table S2.

**Figure 1.**
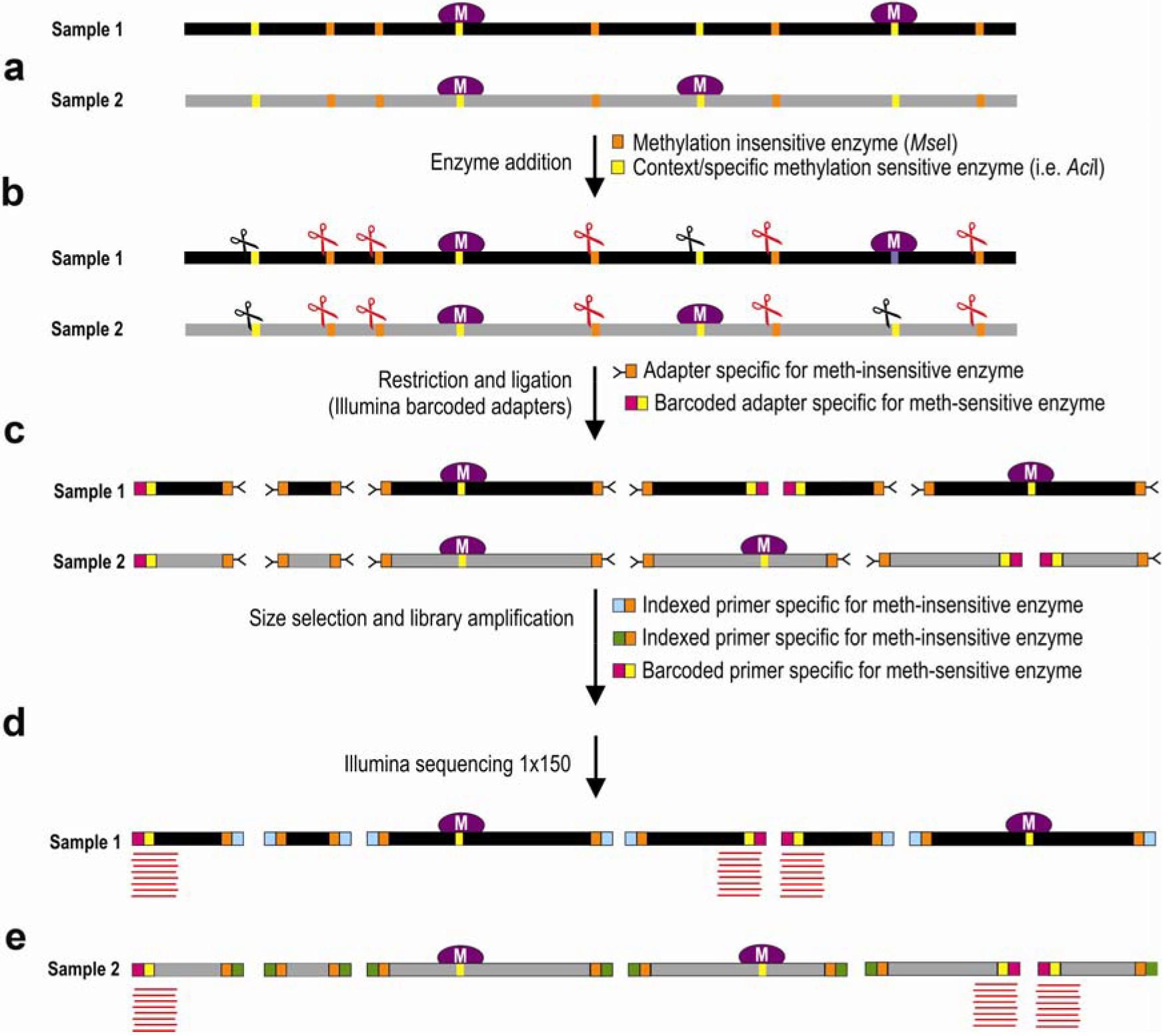
Schematic representation of MCSeEd. Two samples (e.g., control, stressed samples) (a) are subjected to double digestion (b) with a methylation-insensitive (*Mse*I) and a methylation-sensitive enzyme (*Aci*I, *Pst*I, *Eco*T22, *Dpn*II; see Supplementary Table S1) and ligated (c) with specific adapters: a Y adapter for the *Mse*I site and a barcoded adapter specific for the methylation-sensitive enzyme (c). After size selection, the fragments are pooled and amplified with adapter-specific primers (*Mse*I primers carry library-specific indices) and sequenced using the 1×150 Illumina chemistry (d). Demultiplexed reads (e) are analyzed with the design-appropriate MCSeEd pipeline.

A mean of 7.8 million 150-bp-long reads were obtained from each library (Supplementary Table S3). Of these, 98% passed quality control and were aligned to the B73 reference genome. To avoid bias due to paralogous sequences that can align at multiple genomic sites, only the reads that mapped at unique genomic positions were retained. Thus, a total of 89,935,677 reads were mapped uniquely on the reference genome (48.49% of the total reads, with a minimum of 30.13% for *Dpn*II, and a maximum of 78.70% for *Pst*I). We named these reads the MCSeEd loci (Supplementary Table S3). They identified 992,320 loci containing cytosines (705,341 in symmetric, and 286,889 in asymmetric contexts) and 1,629,894 loci containing adenines (Supplementary Table S4).

The mapping location of each MCSeEd locus was investigated to determine whether it fell into a gene window that included the region within 2.5 kb upstream of the transcription start site (TSS), the transcribed region (i.e., the gene body), and the region within 2.5 kb downstream of the transcription termination site (TTS). Of note, in all, 92.9% (*Aci*I), 78.0% (*Pst*I), 82.3% (*Eco*T22I), and 98.1% (*Dpn*II) of the identified MCSeEd loci fell into these gene windows. Specifically, a mean of 6.28 sites per gene window was recorded, with a minimum of 2.96 for *Pst*I, and a maximum of 12.58 for *Dpn*II (Supplementary Table S5).

After normalization of the MCSeEd loci, the sites covered by a total number of reads <4 or showing excessive read-count variation among the replicates (standard deviation >8%) were discarded (Supplementary Table S4). The remaining sites were used to estimate the total of 62,489 DMPs, out of the 992,230 investigated cytosines, with significantly altered methylation levels between the WW and DS samples (false discovery rate, ≤0.05). Of these, 44,176 belonged to symmetric, and 18,313 to asymmetric contexts. With similar filtering, out of 1.6 million on 6mA, 118,269 DMPs were detected (Supplementary Tables S6, S7).

Principal component analysis was used to cluster the samples based on the methylation levels of the DMPs (Supplementary Figure S1). The first latent component (PC1) accounted for 53%, 84%, 44%, and 47% of the total variance, for the CG, CHG, CHH, and 6mA contexts, respectively, and clearly discriminated between WW and DS, which indicated that the drought stress leads to genome-wide methylation changes in maize leaves.

Accordingly, complete linkage clustering of the methylation levels at DMPs clearly separated the DS from the WW samples (Supplementary Figure S2). Altogether, these data indicate that the MCSeEd pipeline can infer the effects of drought stress for each methylated context. Considering all of the methylation changes as being induced by water deficiency in the drought-stressed replicates, we observed 1.6-fold (CG) to 3.4-fold (CHH) more methylation increases than decreases as responses to this stress, whereas for CHG and 6mA, the proportion of methylation changes in each direction were effectively equivalent (Supplementary Figure S3a).

#### Differentially methylated regions

Genomic regions with co-regulated methylation changes upon drought-stress were identified by an adjacent window approach that targeted adjacent DMPs with concordant methylation changes (at least 2). After validation by logistic regression, the identified genomic regions were investigated as DMRs. In total, 5,726 DMRs were identified for the CG (347), CHG (836), CHH (205), and mA (4,338) contexts (Supplementary Table S6). The DMR median length was similar for CG (485 bp), CHH (484 bp), and 6mA (506 bp), and a little lower for CHG (325 bp) (Supplementary Table S8).

The estimated relative methylation level of the DMPs belonging to each DMR were hierarchically clustered, and as expected, clustered according to treatment, as WW or DS (Figure 2). In particular, for the CG and CHH sites, the number of DMRs with higher methylation levels in the DS samples (relative to the WW samples) was higher than the number of DMRs that showed a lower level in the DS samples (Figure 2). In contrast, for the CHG and 6mA contexts, the number of DMRs with higher methylation levels in the DS samples (relative to the WW samples) was equivalent to the number of DMRs with a lower level in the DS samples.

**Figure 2.**
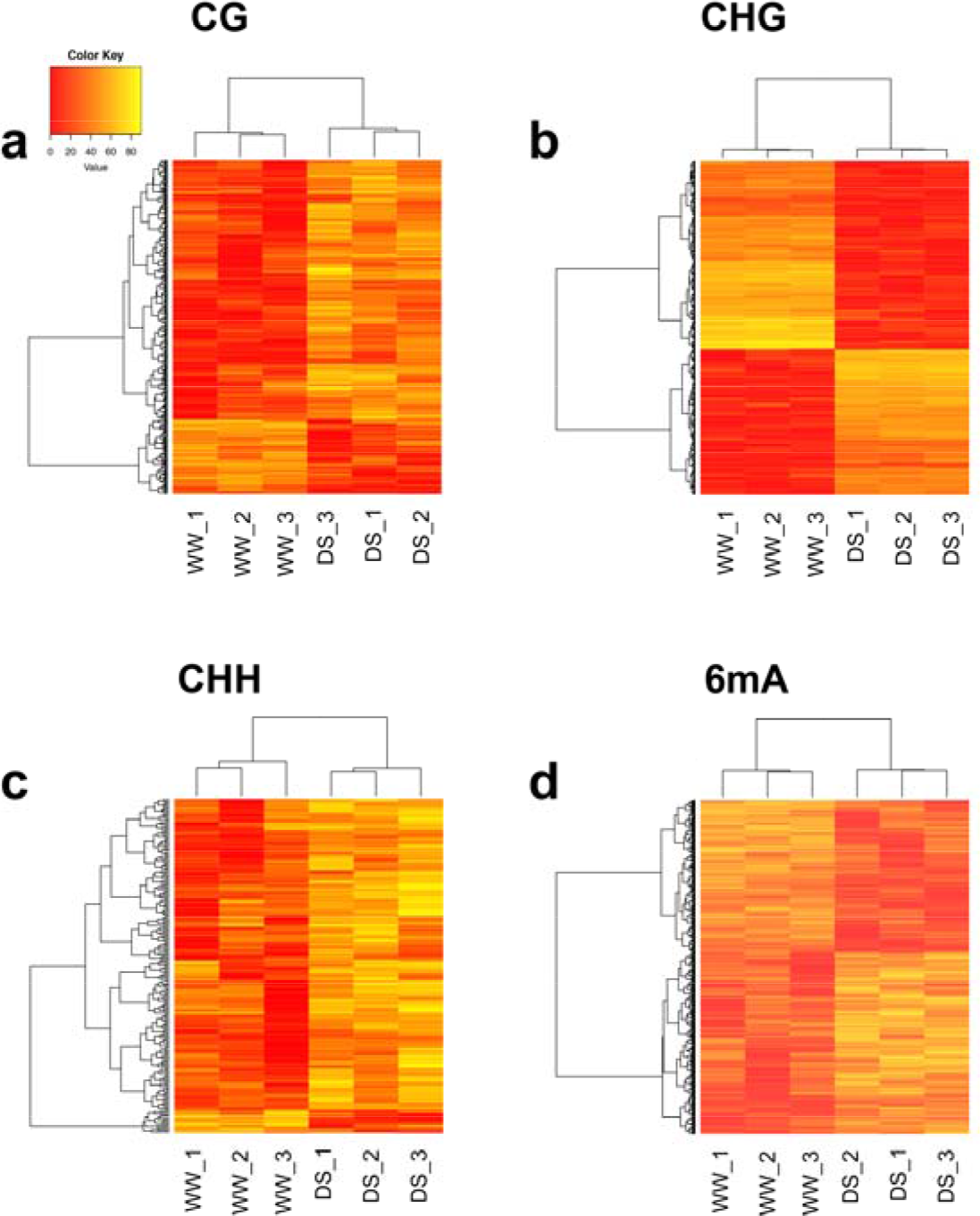
Relative methylation frequencies of differentially methylated regions identified from the comparison between the well watered (WW) and drought stressed (DS) samples. Relative methylation frequencies of the differentially methylated positions contained in each differentially methylated region were averaged and used in complete linkage clustering analysis of samples derived from WW and DS based on 347 (a; CG), 836 (b; CHG), 205 (c; CHH) and 4338 (d; 6mA) differentially methylated regions.

#### Differentially methylated genes

To analyze how water stress impacts the methylation patterns typical of genic regions, we analyzed the DMP and DMR distributions in relation to the coding and regulatory genomic sequences. In particular, we compared the distribution of DMPs and DMRs in transcribed genic regions extended by 2 kb at both ends (extended gene bodies; EGBs) (Supplementary Figure S4, Figure 3). For the DMRs (Figure 3), for all the contexts, they mapped specifically to the gene bodies, and within the 2 kb window upstream of the TSS and downstream of the TTS, while they were depleted in the intergenic regions (Figure 3).

**Figure 3.**
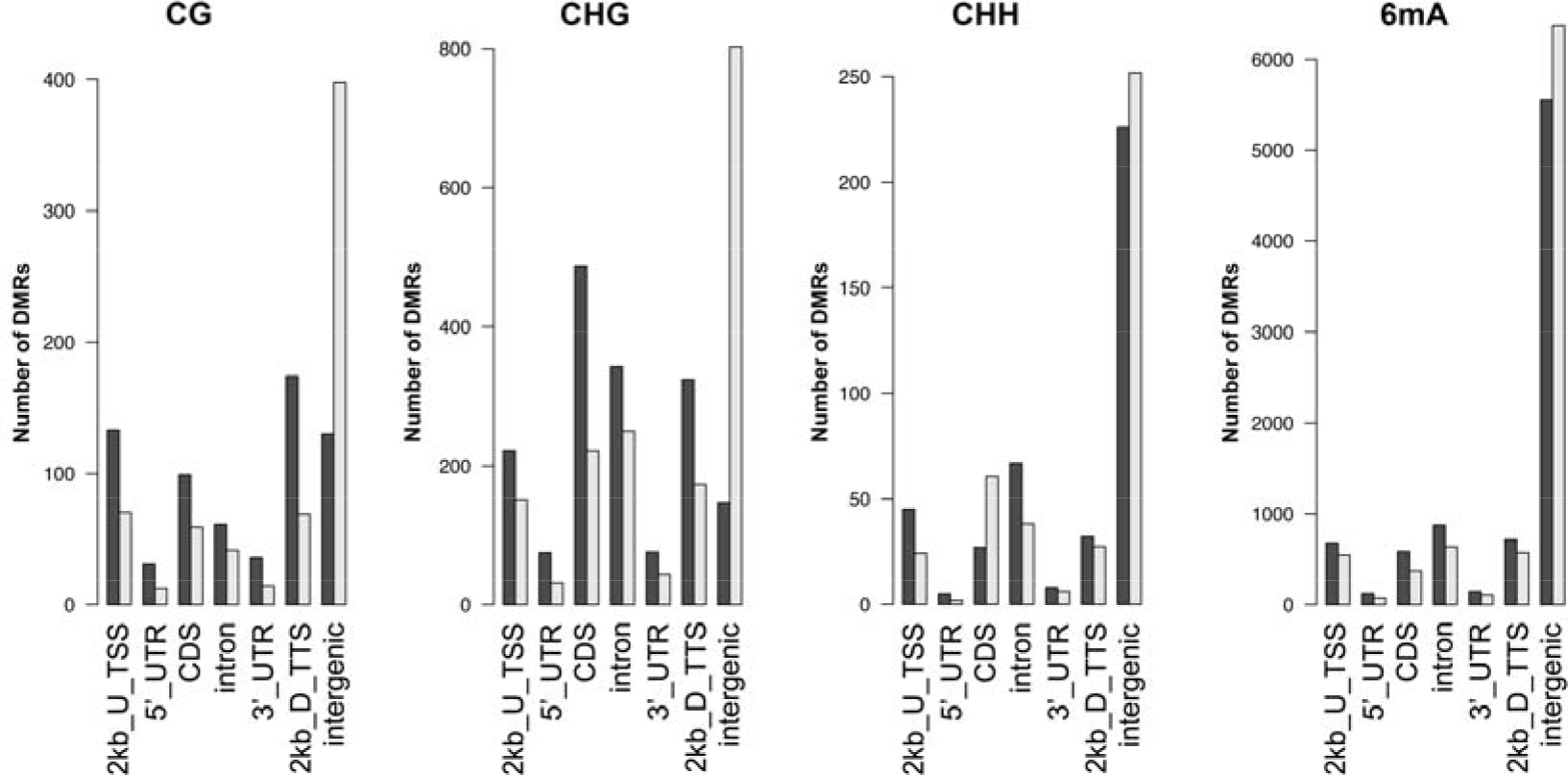
Enrichment analysis of differentially methylated regions in the genomic regions. Enrichment analysis was performed using the binomial distribution of all of the MCSeEd loci as expected and the differentially methylated regions (CG, CHG, CHH, 6mA contexts; note that scales for each context differ), as the observed datasets. U, upstream; D, downstream.

Here, 245 (17.6%), 743 (53.5%), and 307 (22.1%) differentially methylated genes (DMGs) overlapped at least one of the 1,388 DMRs (for CG, CHG, CHH; Supplementary Table S9) within the 2 kb windows upstream of TSS, within the gene body, and within the 2 kb windows downstream of TTS, respectively. These loci were investigated as DMGs upon drought stress. Moreover, 418 (9.6%), 933 (21.5%), and 448 (10.3%) DMGs overlapped at least one of the 4,338 6mA DMRs within 2 kb upstream of TSS, within the gene body, and within 2 kb 2 kb downstream of TTS, respectively.

Panther enrichment analysis using all of the DMGs identified in all of the contexts identified the gene ontology (GO) terms, which were mainly related to regulation of transcription, biosynthetic and metabolic processes, responses to stimuli, oxidoreductase activity, and binding of nucleic acids (Supplementary Table S10). Several DMGs identified from DMRs located within 2 kb upstream of the TSS have orthologs in other species (i.e., *Arabidopsis*) including DRE-binding protein 3 (Zm00001d021207), WRKY 36 (Zm00001d039532), ethylene-responsive transcription factor WIN1 (Zm00001d046501), Myb family transcription factor PHL6 (Zm00001d015226), ISWI chromatin-remodeling complex ATPase CHR11 (Zm00001d040831), with role related to response to water deprivation, defense responses, drought stress tolerance, dehydration stress memory, and response to drought stress, respectively (Supplementary Table S11).

### Validation of differentially methylated positions inferred by MCSeEd

The MCSeEd technique was validated using the two most common methods for DNA methylation analysis: (i) quantitative (q)MRE^37–40^; and (ii) bisulfite cytosine conversion. In the first technique, if the cytosine of the restriction site within a PCR-amplification target is methylated, the enzyme cannot cut the DNA, and the relative amplicon is produced; conversely, when the cytosine is not methylated, the DNA is digested by the enzyme and the amplificon cannot occur. The second technique, determines if a cytosine is methylated or not via bisulfite conversion of DNA, whereby unmethylated cytosines are converted into thymidine, while methylated cytosines remain unchanged.

For the MRE approach, 10 randomly chosen DMPs were used. Nine of these 10 DMPs showed methylation differences comparable to those obtained after methylKit analysis (Supplementary Table S12), while the remaining one showed no significant differences between the WW and DS samples.

For the second comparative analysis, MCSeEd was applied to shoots and roots of the B73 maize inbred line collected 5 DAS (Supplementary Table S3). Data retrieved from the MCSeEd shoot samples were then compared with two public datasets of shoot-WGBS as benchmark data^41^, according to Maunakea et al.^29^. In particular, the MCSeEd scores (normalized number of reads interrogating cytosines for the CG, CHG, and CHH contexts) were compared with bisulfite sequencing scores (number of reads for the methylation content for the CG, CHG, and CHH contexts). These data showed negative correlation for the CG context (Spearman correlation, −0.506; *P*-value, 2.2 e-16) and the CHG context (Spearman correlation, −0.517; *P*-value, 2.2 e-16; Figure 4). For CHH, no correlation was seen (Spearman correlation, 0.012; *P*-value, 1.1 e-09; Figure 4), probably due to the lower number of read counts.

**Figure 4.**
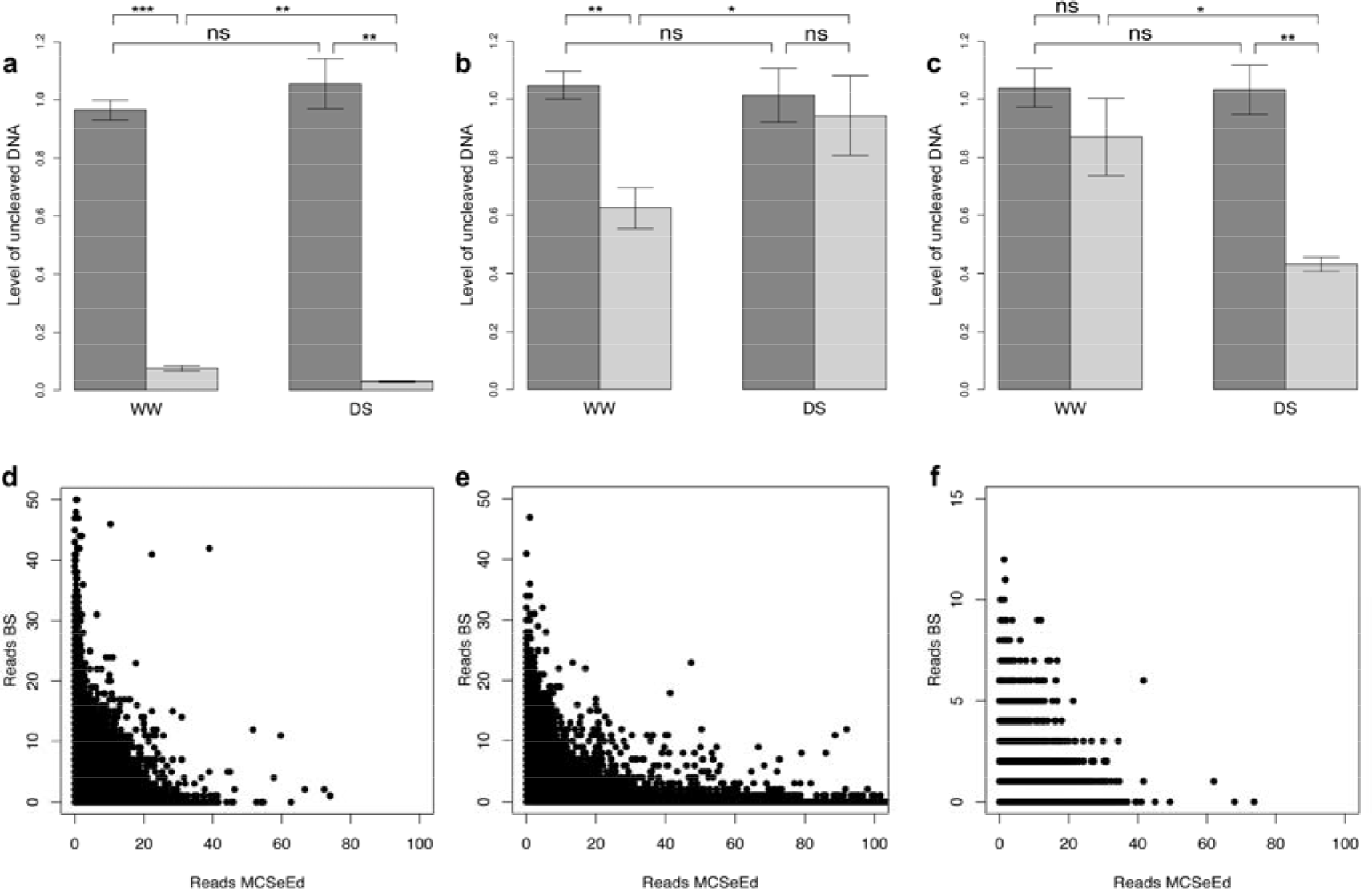
The MCSeEd validation by quantitative methylation-sensitive PCR (a-c) and comparisons with WGBS (d-f), for the well watered (WW) and drought stressed (DS) samples. (a) DMP_X. (b) DMP_Y. (c) DMP_Z (Supplementary Table S12). Dark shading, mock (no digested DNA, considered fully methylated); light shading, digested DNA. Error bars indicate standard deviation. *, p <0.05; **, p <0.01; ***, p <0.001; ns, p >0.05 (unpaired two-tailed Student’s t-tests). Spearman correlation test between WGBS datasets^41^ and MCSeEd loci for CG (d), CHG (e), and CHH (f). Unmethylated cytosines are shown as MCSeEd reads (normalized number of reads interrogating each cytosine) on the X-axis. Methylated cytosines are shown as bisulfite sequencing reads on the Y-axis.

### Reference-free strategy

Since one of our goals was to develop a high-throughput technique that can also be applied to species without a reference genome, the MCSeEd bioinformatic pipeline was tested for mapping of filtered reads to a pseudo-reference genome autogenerated by a pipeline, hereafter indicated as the genome independent strategy.

Under the DS conditions, a total of 2,258,361 pseudo-genome contigs were generated by methylation content (686,548 contigs, 30.4%) and 6mA (1,571,813, 69.6%) libraries (Supplementary Table S4, methylKit). After MCSeEd loci normalization, the sites covered by a total number of reads either <4 or showing an excessive read count variation among the replicates (standard deviation >8%) were discarded (Supplementary Table S4), and the remaining sites were used to estimate the total of 30,092 DMPs, out of 686,548 investigated cytosines, with significantly altered methylation levels between the WW and DS samples. Of these, 27,203 belonged to symmetric and 10,889 to asymmetric contexts. Moreover, out of 1.6 million on 6mA, 143,389 DMPs were detected (Supplementary Tables S6, S7).

In the genome-independent approach, both principal component analysis and heatmaps (Supplementary Figures S5, S6) clearly discriminated between the WW and DS samples. The CG and CHH contexts showed higher methylation in response to stress (2.89-fold, 5.09-fold, respectively), while for CHG and 6mA, the differences there were effectively no differences (0.92, 0.88, respectively; Supplementary Figure S3b).

### Variant calling and shift in allelic frequency

Allele-specific methylation responsive (ASMR) sites to drought stress were identified based on the allelic origins of the reads. Using SNP information from *Pst*I libraries as an example, the relative allelic contributions to the total read counts were inferred by counting the number of reads that originated from either SNP allele (Supplementary Table S13). This highlighted 10,861 heterozygous SNP loci. Among these, 287 showed significant shifts in their relative allelic contributions between the WW and DS samples (P <0.05), and were defined as ASMR sites (Figure 5).

**Figure 5.**
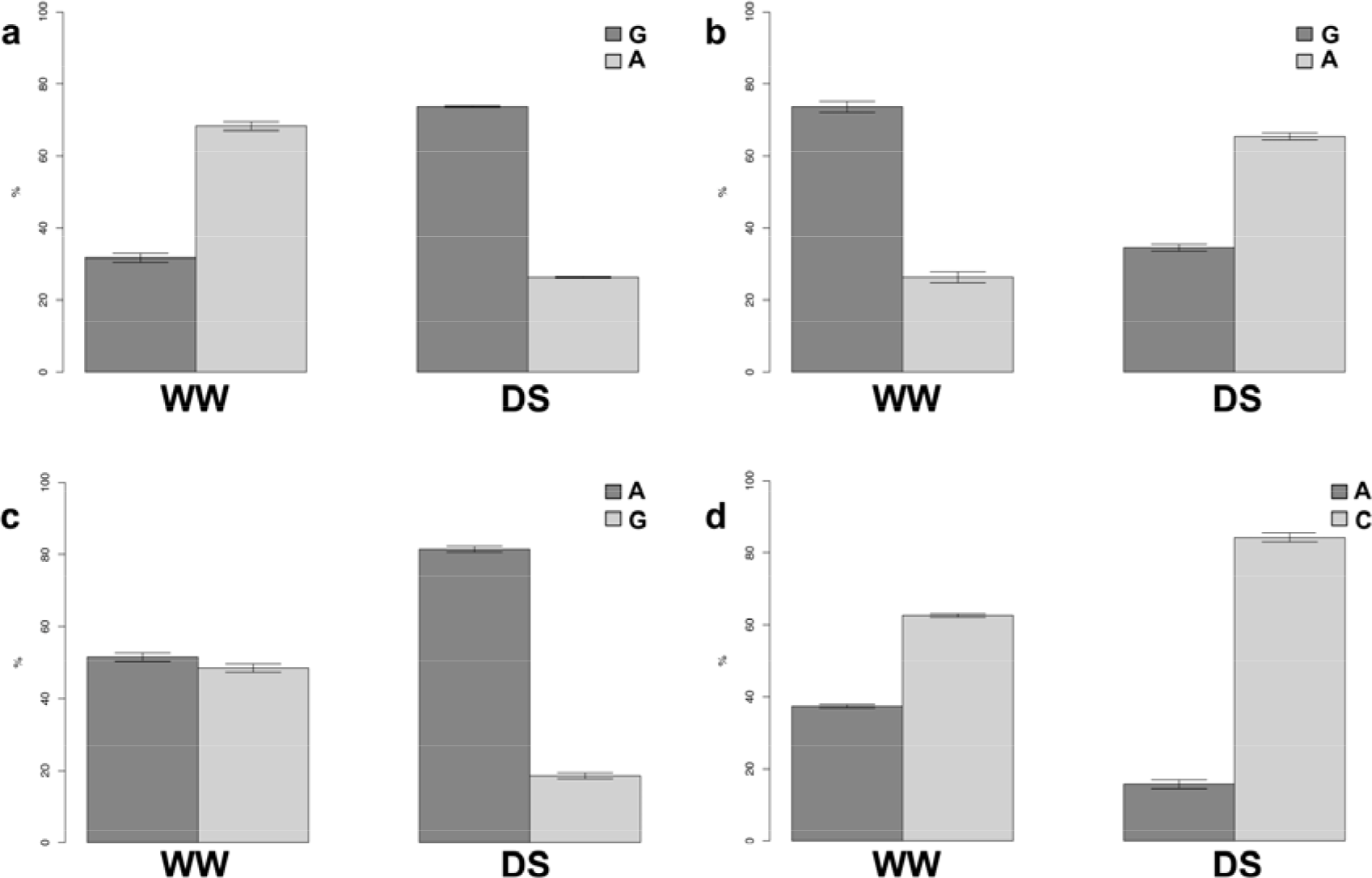
Examples of the differential changes in the methylation states due to drought stress, between the alleles at the single SNP loci. *Pst*I libraries were screened for SNP polymorphisms at each of the four random loci: (a) Chr1_254513757; (b) Chr1_34649135; (c) Chr8_150564756; and (d) Chr8_169547803. WW, well watered; DS, drought stressed.

A total of 128 of the 287 identified ASMR sites also showed significant net changes in total methylation status (i.e., the DMP–ASMR loci). The remaining 159 ASMR loci were not related to DMPs. This might have been because the methylation changes between WW and DS for one allele was compensated for by a similar change in the opposite direction for the other allele, therein not altering the total relative methylation levels between WW and DS. Interestingly, the ASMR sites were preferentially located (223 out of 287) in EGBs (P <0.05): 128 within genes, 52 within 2 kb upstream of the TSS, and 43 within 2 kb downstream of the TTS (Supplementary Table S14).

## Discussion

We have developed a reduced-representation, reference-free approach for characterizing whole genome methylation patterns across different methylation contexts. While WGBS is the gold standard, it is cost prohibitive at most experimental scales, and requires a reference genome (Young et al. 2016).

Alternative approaches aimed at reducing the sequencing demands are based on genomic digestion with either methylation-sensitive or methylation-insensitive endonucleases, before direct NGS (e.g., EpiRADseq^30^, MRE-seq^29^), or bisulfite treatments and NGS (e.g., RRBS^27^, methylation-sensitive restriction enzyme bisulfite sequencing [MREBS]^42^). However, both RRBS and MREBS still require a reference genome and a certain level of coverage, at least for estimation of the methylation status of the cytosines. Although EpiRADseq and MRE-seq do not require a reference genome, these consider only the CG sites, and the methylation status is deduced by counting the unmethylated cytosines within the recognition site. The scenario for 6mA is even worse to date, with the level of 6mA determined by immunoprecipitation or liquid chromatography coupled with tandem mass spectrometry^7,32^. The only sequencing technique that can simultaneously infer the cytosine-5 methylation and 6mA is PacBio single-molecule real-time sequencing, but this is prohibitively expensive for most applications^7,32^.

To address these limitations, we developed a new method: MCSeEd. Similar to other reduced-representation methods, MCSeEd is based upon parallel restrictions carried out by combinations of a methylation-insensitive endonuclease (*Mse*I) and one of four methylation-sensitive endonucleases, directed to one of CG, CHG, CHH, and 6mA. This is completed using NGS.

With MCSeEd, the read counts can be readily used to estimate the differential methylation between two samples, which is often of primary interest^42^. Indeed, the cytosine libraries here described were equally represented, and they covered a mean of 2.6 million sites, about 7% of which were differentially methylated between the samples. This MCSeEd pipeline was tested without and with the support of a reference genome and in both cases, this resulted in identification of DMPs at very high rates. The genome-dependent approach resulted in a similar number of DMPs with respect to the genome-independent approach (180,758 vs 181,481).

We used two validation systems. The first was qMRE, which was used to confirm the changes in the DNA methylation recorded in the WW *versus* DS comparison. The second was the comparison between the two datasets, with shoot WGBS as the benchmark data^41^, and the MCSeEd data obtained at the same stage for the same inbred line (B73). To confirm our DS experiment data, we randomly chose 10 DMPs and applied the qMRE method. Of note, this system was able to confirm the methylation levels within the samples and the differential methylation between the WW and DS samples in nine out of the 10 cases.

In the comparative analysis following the MCSeEd validation, negative correlations were seen for the comparison of these MCSeEd GC scores and the WGBS CG and CHG scores (−0.506, −0.517, respectively). These data are similar to those of Maunakea et al.^29^ for MRE-seq, and they support the idea that MCSeEd is focused on genome sequences that are most likely unmethylated, and relies on the inverse relationship between read coverage and DNA methylation level at a given locus. When we compared MCSeEd and bisulfite sequencing data for the CHH context, there were no correlations. A possible explanation here is that the CHH sites are scarcely methylated, and the low cover of bisulfite sequencing fails to identify these positions as methylated. Indeed, 93% of the genomic positions identified with MCSeEd had 0 reads in the bisulfite sequencing dataset. On this basis, both MRE-seq and MCSeEd are effective in revealing loci with low frequencies of *de-novo* methylation that are generally missed by the WGBS and RRBS methods (unless performed at very high coverage), which makes MCSeEd one of the techniques of choice when looking for comparative changes in methylation status related to gene expression.

Methylation of adenines is still a poorly investigated field. For instance, Fu et al.^14^ reported that the consensus motifs for 6mA are only partially conserved in different eukaryotic organisms, and enzymes might have evolved to catalyze 6mA modifications in the evolutionary process. They reported that GATC appears to be the most ancient 6mA motif, which exists in both lower eukaryotes and bacteria, but is lost in higher eukaryotes^14^. Instead, GAGG is present in both *plantae* and *animalia*, and AGAA might be specific to animals^15,43^. Liang et al.^7^ demonstrated that *Arabidopsis* contains two specific methylated motifs, ANYGA and ACCT, which have not been found in other organisms to date. On the other hand, the presence of adenine methylation at GATC sequences was shown in rice^44^, tobacco^45^, and *Arabidopsis*^46^. In particular, both Dhar et al.^44^ and Ashapkin et al.^46^ used *Dpn*I, which digests the DNA only if adenine is methylated in the sequence GATC, to demonstrate the extensive digestion of rice and *Arabidopsis* DNAs. Our results confirm these findings, and demonstrate that methylation at the GATC site is not only present in maize, but also that the mechanisms of methylation/ demethylation are still active, as the levels of differentially methylated positions due to stress are very high (118,269 and 143,389 DMPs in the genome-dependent and genome-independent approaches).

CG methylation is prevalent in the transcribed regions of many constitutively expressed plant genes (i.e., gene body methylation), and it shows characteristic patterns within genic regions^47,48^. In maize, CG methylation has been shown for exons more than introns, which suggests a defensive function *versus* transposon insertion in the coding sequence, while allowing insertions into introns and other noncoding regions^41^. Moreover, both in maize and *Arabidopsis*, the CHG and CHH methylation contexts are significantly enriched at intron-exon junctions^41^. Our data show that all cytosine-5 methylation contexts are enriched within 2 kb upstream of the TSS; moreover, while CG and CHG are highly enriched in exons, CHH is enriched in introns.

For 6mA, Liang et al.^7^ reported that these sites in *Arabidopsis* genes are mainly located in exons, while those in transposable element genes show a local reduction at the TSS, followed by an immediate increase, as also seen in *Chlamydomonas* genes^14^. This is in contrast to what was seen in *C. elegans*, where 6mA are distributed equally in genomic regions, including introns, exons, and TSS regions, and in *Mus musculus* and *Xenopus laevis*, where 6mA are primarily excluded from coding regions. Our data show an enrichment of 6mA in EGBs, and lower levels of 6mA in intergenic regions. This is particular true for DMRs, but less evident for DMPs. It is worth nothing that the 6mA located in EGBs appears to respond to water restriction more than the 6mA located in intergenic regions.

The efficiency of MCSeEd for detection of changes in DNA methylation between two sets of contrasting samples was evident in both principal component analysis and linkage clustering, which clearly discriminated the WW samples from the DS samples, for both DMPs and DMRs.

Boyko et al.^49^ reported an increase in global genome methylation in *Arabidopsis* plants exposed to stress, including salt, UVC, cold, heat and flood stresses. In particular, the progenies of these stress-treated plants showed increased global methylation, even in the absence of the stress, but these transgenerational effects did not persist in successive generations in the absence of stress. Moreover, induction of transient DNA methylation was related to drought stress in pea^50^ and in drought susceptible rice genotypes^51^. By using MCSeEd on these maize samples with different water status (i.e., WW vs. DS), we found that DS appears to induce methylation rather than demethylation, particularly in the CHH and CG contexts. In total, we identified 1240 DMGs, most of which were related to regulation of transcription, biosynthetic and metabolic processes, and response to stimuli, processes that suggest that changes in DNA methylation are correlated with stress responses to water deprivation.

Genomic DNA cytosine/ adenine methylation polymorphism studies can be tested for their ability to reveal “epigenetic heterosis” effects for tolerance to drought and other stress factors, and as a tool to identify novel stress-responsive genes. Indeed, methylation changes can be allele specific or genome specific (in case of polyploids). Therefore, inferred ASMR sites might be very important for this purpose. In our study, many ASMR sites for drought stress were identified, as a genome-wide situation that already existed in the control plants (i.e., WW). It is known than genomes can show selective allelic imbalance, as seen for humans^52^ and plants^53–55^.

The drought stress influenced the methylation status of cytosines and adenines, which created a level of imbalanced heterozygosis between the stress (DS) and control (WW) conditions. This imbalance might impact on the level of gene expression, as methylation can have a general role in the regulation of gene expression and contribute to the hybrid vigor phenomenon^56–59^. MCSeEd can thus mine SNPs associated with selectively methylated sites, to highlight allelic imbalance. This has implications for breeding programs, wherein MCSeEd can provide information about which of two genomes in a hybrid has been methylated, and through the targeting of candidate ASMR sites, allow a “MAS by methylation” approach.

Here we tested MCSeEd on maize, as an important crop that has a large genome and is thus very rich in transposable elements. Also, in its cultivated form, maize is a hybrid, and hence a very challenging species to work with. We have demonstrated that even in such a complex scenario, use of MCSeEd identified differentially methylated regions both between stress and control conditions in a hybrid genotype, and between developing organs in an inbred line (B73). Benchmarking experiments indicate that MCSeEd can be reliably used for differential methylation analyses with consistent results.

## Materials and methods

### Plant material

For the drought-stress study, plants of a commercial maize hybrid variety were subjected to normal irrigation (well-watered; WW) and drought stress (DS), as reported by Bocchini et al.^60^. At 60 DAS, portions of the leaves were collected, bulked into three biological replicas of five plants each (WW1, WW2, WW3, DS1, DS2, DS3), and then stored at −80 °C, until further processing.

For MCSeEd validation, B73 seeds were germinated at 25 °C in the dark on wet paper towels in glass Pyrex dishes. At 5 days after germination, the shoots at the coleoptile stage and the roots were excised and stored at −80 °C, until further processing.

### DNA purification, library construction and sequencing

Genomic DNA was purified from each sample using DNeasy Plant Mini kits (Qiagen GmbH, old (Qubit; Life Technologies, Grand Island, NY, USA). The library set-up protocol was performed according to Peterson et al.^31^ with some modifications, as described below. Four specific enzyme combinations were chosen (as one of four methylation-sensitive enzymes, each combined with methylation-insensitive *Mse*I) to infer the CG (*Aci*I/*Mse*I), CHG (*Pst*I/*Mse*I), CHH (*Eco*T22I/*Mse*I), and 6mA (*Dpn*II/*Mse*I) methylation contexts, respectively (Supplementary Table S15, Figure 1). To define the efficacy of the enzyme combinations, we developed a program that scanned the genome *in silico* and calculated the size distribution of the restriction fragments. Briefly, the user was asked to insert the name of a desired restriction enzyme combination, and a virtual digestion was performed. Fragments with the optimal length range can then be selected and counted.

For each library, 150 ng DNA were double-digested with one of these four enzyme combinations. In the same reaction, a sample-specific barcoded adapter was ligated to the methylation-sensitive restriction end, while a common Y adapter was ligated to the sticky end left by *Mse*I (Supplementary Table S2). The libraries were then pooled, as reported in the experimental design (Supplementary Table S2), purified using magnetic beads (Agencourt AMPure XP; Beckman Coulter, MA, USA), size selected by gel electrophoresis, and purified using QIAquick Gel Extraction kits (Qiagen) for fragments in the range of 200 bp to 650 bp. Size-selected libraries were quantified using a fluorometer (Qubit; Life Technologies), and a normalized DNA amount was amplified with a primer that introduced an Illumina index for demultiplexing. Following PCR with uniquely indexed primers, multiple samples were pooled. PCR-enrichment was performed as described by Peterson et al.^31^. Amplified libraries were purified with magnetic beads (AMPure; Beckman Coulter, Brea, CA, USA), and then quantified (Qubit and Bioanalyzer 2100: Agilent Technologies, Santa Cruz. CA, USA). The grouped libraries were pooled in an equimolar fashion, and the final library was Illumina-sequenced using 150-bp single-end chemistry.

### Genome-dependent workflow

Raw reads from the Illumina sequencing of the CG, CHG, CHH and adenine methylation libraries were demultiplexed using the *process_radtags* tool (STACKS v.2.3b package)^61^. This identifies and assigns reads to each individual on the basis of 7-bp custom barcode sequences (removed after analysis). After processing the raw reads, the MCSeEd pipeline was run following either genome-dependent or genome-independent procedures, as detailed below. The MCSeEd pipeline consisted of a bash wrapper using different algorithms or Perl scripts. Sequences from each library were mapped to the reference maize genome (AGPv4; https://www.maizegdb.org) with the *bwa mem* algorithm using the default settings^62^. Bam sorted and indexed files of uniquely mapped reads were produced with *Samtools*^63^.

### Loci-counting approach

To create a count matrix where the columns are the sample libraries and the rows represent the locations in the genome hit by the sequencing, we created a merged bam file that acted as a guide for creating an “experiment-wise annotation”. Here, all of the genomic positions sequenced were stored, whereby all of the uniquely mapped reads were recorded for each genomic location covered in the experiment. Briefly, for each alignment in the bam file, genome coordinates and CIGAR fields were processed to produce meaningful intervals with a Perl script. Then, redundant coordinates were collapsed and sorted, and overlapping intervals were merged with the *bedtools* suite, maintaining strandedness^64^. This information was converted to a formal GFF file, and then used as the input in *featureCounts*^65^, along with the bam files previously described, to count the occurrences on the experiment-wise annotation.

The count matrix consisted of one row per locus and one column per sample, and it was then filtered and processed by two in-house–built Perl scripts. The following operations were performed: (1) the libraries were normalized and balanced in a reads per million fashion; (2) loci with a coverage of at least 10 reads were retained; (3) relative methylation level per locus were estimated; and (4) the filtered data were parsed for use by the methylKit R package^66^. In particular, the relative methylation levels at each site (point 3) were calculated following a rescaling procedure that was based on the maximum number of observed read counts. In practice, the sample showing the highest number of reads was assumed to be the not-methylated reference for the site, or to have 0% methylcytosine and 100% cytosine, and all of the remaining samples were rescaled proportionally (Supplementary Table S16). As the reference was common to all samples, the methylation level estimates can be used to infer relative methylation changes between the samples. DMPs were therefore identified as sites that showed significant differences in the methylation levels between the treatments, using logistic regression as implemented in methylKit. The DMPs were called following the methylKit manual best practices.

The mapping of the DMPs in the same scaffold and as closer than a given threshold provided their clustering together to identify the DMRs, based on the following procedure. Briefly, the first step was to maximize the number of DMRs in a set of adjacent windows, to identify the best window length for each context. We therefore tested a range of windows, from 100 bp to 2000 bp. To do so, each potential window (i.e., 100 bp) was screened for DMPs that were significantly differentially methylated (false discovery rate, ≤0.05). The 5’-end of the window was therefore registered to start at the DMP position. Additional DMPs that were mapped within the re-positioned window (i.e., 100 bp) were included in the cluster, provided that the following conditions were met: (i) the direction of the methylation change agreed with the preceding DMP included in the cluster; and (ii) the DMPs to be included were called with a given significance threshold (false discovery rate, ≤0.05). After the additional DMPs were included in the cluster, the window start was registered to the position of the most 3’ of the DMPs included, and the procedure was repeated as described. If no additional DMPs were identified based on the described condition, the scanning procedure was restarted until a DMP was identified. These clusters of DMPs that were composed of a number of DMPs that exceeded a given threshold were analyzed using logistic regression, to identify and define the DMRs.

Once the data for each window length was produced, the operator chose the best length, i.e., the one that maximized the number of DMRs per window (Supplementary Table S8). At this point, the script was re-stared for each context using the adjacent window of the chosen length.

### Genome-free workflow

For the genome-independent part of the pipeline, we relied on the robust and simple approach of Schield et al.^30^, with some modifications. Briefly, as no reference genome was available, the raw reads were collapsed using Rainbow 2.0.4 (https://sourceforge.net/projects/bio-rainbow)^67^ and CDHit (https://github.com/weizhongli/cdhit)^68^, to create a pseudo-reference genome that consisted of a multi-fasta file that contained the read contigs: this serves as a guide for the mapping algorithm. After mapping the reads to the pseudo-reference using the *bwa mem* algorithm with its default settings^62^, a result matrix was created for each sample using Samtools, which counts how many sequences per contig are mapped. This matrix was then used following the loci counting approach described above.

### Variant calling and unbalanced allelic frequency mining

For both genome-dependent and genome-independent approaches, MCSeEd can perform a variant calling procedure using the Stacks suite^61,69^. Briefly, after the creation of a population file and a catalog of mutations, a VCF file with frequencies and reference/ alternative alleles was created, which was ready to be transformed into any population-genetics exchange file (i.e., PED/MAP).

To highlight unbalanced allelic frequencies in heterozygous loci among the WW and DS samples, and putatively due to differential methylation, the reference (REF) and alternative (ALT) allele counts were extracted from the vcf files and normalized across the samples. Only sites covered by at least 25 reads were considered. Then, at each site, the frequency of the reference allele was computed for each allele across the three replicates, as in Equation (1):

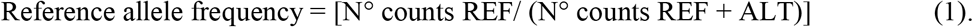

This was then used to calculate the mean frequency at each locus, for each enzymatic context. The shifts in the allele contribution indices between the WW and DS samples were calculated based on Student’s t-tests, followed by Benjamini-Hochberg correction for multiple tests (P <0.05).

### MCSeEd validation using qMRE

As described by Hashimoto et al. (2007), for each MCSeEd sequence to be validated, the DNA was digested using the methylation-sensitive enzyme for the corresponding methylcytosine context. The reaction mixture of 25 μL contained 100 ng DNA, 0.5 U of either *Aci*I, *Pst*I, or *Eco*T22I in the specific buffer defined for each enzyme. For the non-enzyme control (mock), distilled water was added instead of the enzyme. All of the samples were then incubated at 37 °C for 4 h, follow by heat inactivation at 65 °C / 80 °C for 20 min.

Real-time PCR for the methylation status was performed (Mx3000P QPCR system; Stratagene, La Jolla, CA, USA) with the SYBR Green JumpStart Taq ReadyMix for Quantitative PCR (Sigma Aldrich). Using the Primer3 software^70^, specific primers were designed for each randomly chosen DMP to be validated. The sequence information of the primers that bracketed the enzyme site of each DMP are reported in Supplementary Table S15.

The PCR fragments were analyzed using a dissociation protocol, to ensure that each amplicon was a single product. The amplicons were also sequenced to verify the specificities of the targets. The amplification efficiency was calculated from the raw data using the LingRegPCR software^71^.

All of the qMRE were performed in a final volume of 25 μL that contained 20 ng DNA template (previously digested/ mock), 0.2 μM of each primer, and 12.5 μL 2× PCR Master Mix, according to the manufacturer instructions. The following thermal cycling profile was used: 95 °C for 10 min, followed by 40 cycles of 95 °C for 10 s, 57 °C for 15 s, and 72 °C for 15 s. Following the cycling, the melting curve was determined in the range of 57 °C to 95 °C, with the temperature increment of 0.01 °C/s. Each reaction was run in triplicate (as technical replicates).

The raw Ct data from the real-time PCR were exported to a data file and analyzed using the GeneEx Pro software^72^. During the pre-processing phase, the data were corrected for PCR efficiency, with the means of the three biological samples calculated. The selected reference gene, GAPDH (GenBank accession no. X15596.1), was subsequently used to normalize the Ct values^73–75^, and the quantities were calculated relative to the maximum Ct value. As our interest was in fold-changes in the amplification between the mock and treated samples and the WW and DS samples, we ultimately converted the quantities to a logarithmic scale using log base 2 conversion, which also allowed the normal distribution of the values to be tested^76^.

### MCSeEd validation using WGBS

The MCSeEd validation was also performed according to Maunakea et al.^29^, by comparing two sets of shoot-WGBS data (reference, GSM958914, GSM958915) with our data for the shoot samples at the 5 DAS stage. In particular, the two WGBS datasets were mapped using the Bismark software (version 0.19; https://www.ncbi.nlm.nih.gov/pubmed/21493656) on the maize reference genome AGPv4 (https://www.maizegdb.org). Spearman correlation was calculated using normalized numbers of reads interrogating the CG, CHG, and CHH contexts (MCSeEd scores), and the number of WGBS reads for the methylcytosine context (bisulfite sequencing scores).

### Statistical analysis

Statistical analyses were performed in R version 3.3.2 (www.r-project.org) using the ‘stats’, ‘factoextra’, and ‘gplots’ packages. The ‘stats’ package was used to estimate correlations and binomial and logistic regression, and ‘factorextra’ was used to perform the principal component analysis. Complete linkage clustering was carried out using the ‘heatmap.2’ function of the ‘gplots’ package, in combination with the ‘hclust’ and ‘dist’ functions, and with ‘ward.D2’ as the clustering method. The MethylKit R package was used to estimate the methylation changes between the WW and DS samples.

### Availability

The MCSeEd suite and ancillary scripts are available online at https://bitbucket.org/capemaster/mcseed/src/master/.

## Acknowledgements

The authors acknowledge David Haak (Virginia Tech, VA, USA), Cristian Forestan (University of Padova, PD, Italy), Daniele Rosellini and all the other people at University of Perugia for the critical revision of the manuscript.

## Author Contributions

G.M. and E.A. devised the study, and funded the experiments. G.M., C.C., E.P., and E.A. developed the technique. G.M., A.P., and S.C., developed the bioinformatic pipeline. G.M. and C.C. performed the library preparations. G.M., A.P., S.C. and A.A. performed the bioinformatic analysis. Investigation. G.M., A.P., E.A. designed and interpreted all the experimental data. G.M., A.P., and E.A. drafted the manuscript. All authors reviewed and edited the manuscript.

All authors read and approved the final manuscript.

## Data availability

All sequencing data that support the findings of this study can be found under the bioproject <u>PRJNA533220</u>

For the validation experiment using B73 data, we downloaded the shoots WGBS data (GEO Series accession number <u>GSM958914</u>, <u>GSM958915</u>) and used the set of data regarding 5 DAS stage

## Code availability

All custom scripts have been made available at available at https://bitbucket.org/capemaster/mcseed/src/master/.

## Supplementary Figure captions

**Figure S1**. Principal component analysis (genome dependent) for the relative methylation levels at the differentially methylated positions obtained across all of the replicates in each sequence context: (a) CG; (b) CHG; (c) CHH; (d) 6mA. Numbers in brackets indicate the fraction of overall variance explained by the respective component (Dim1, Dim2).

**Figure S2.** Complete linkage clustering based on the relative methylation levels at the differentially methylated positions, as retrieved from the genome-dependent strategy.

**Figure S3.** Effects of drought stress on the global methylation: (a) genome dependent; (b) genome independent.

**Figure S4**. Enrichment analyses of the differentially methylated positions in the genomic regions. Enrichment analysis was performed using binomial distribution for all of the MCSeEd loci as expected, and the differentially methylated positions for the CG, CHG, CHH, and 6mA contexts, as the observed datasets. U, upstream; D, downstream.

**Figure S5**. Principal component analysis (genome independent) of the relative methylation levels at the differentially methylated positions obtained across all of the replicates in each sequence context: (a) CG; (b) CHG; (c) CHH; (d) 6mA. Numbers in brackets indicate the fraction of overall variance explained by the respective component (Dim1, Dim2).

**Figure S6.** Complete linkage clustering based on the relative methylation levels at the differentially methylated positions, as retrieved from the genome-independent strategy.

