## Supplementary figures and images for "Methylation content sensitive enzyme ddRAD (MCSeEd): a reference-free, whole genome profiling system to address cytosine/ adenine methylation changes"

### Supplementary Figure 1

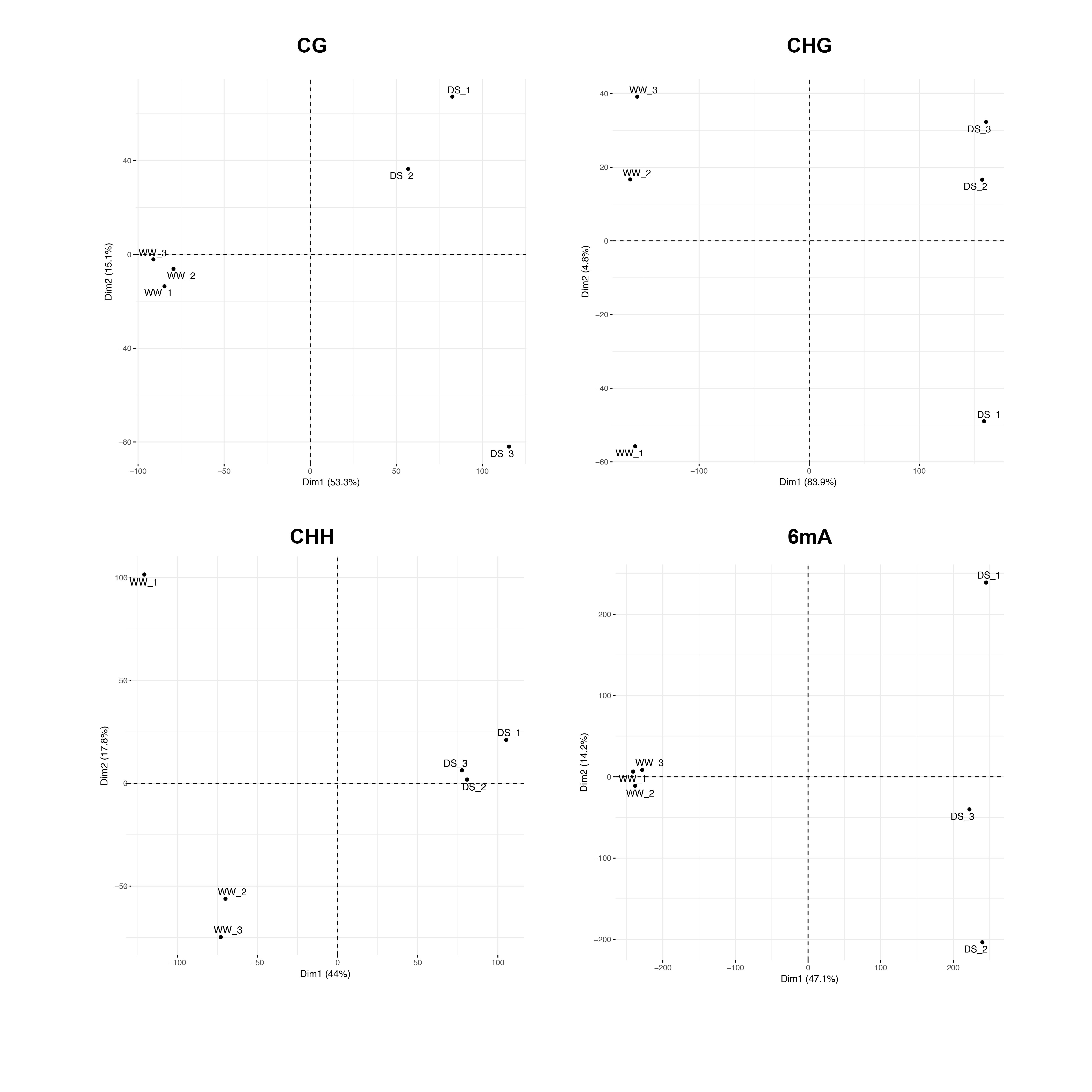

### Supplementary Figure 2

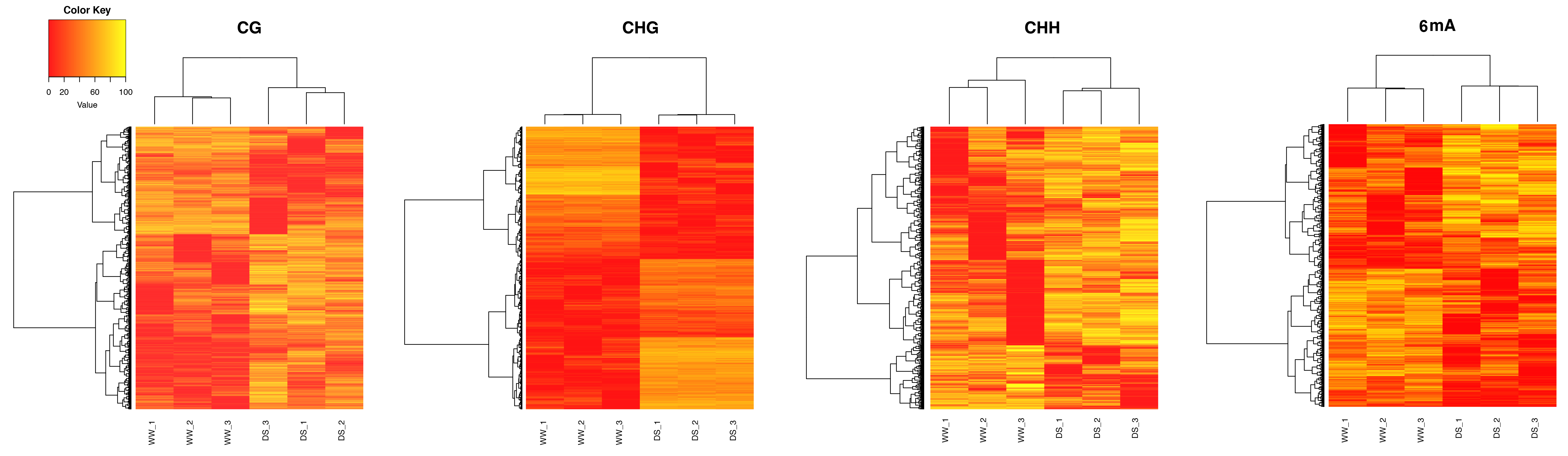

### Supplementary Figure 3

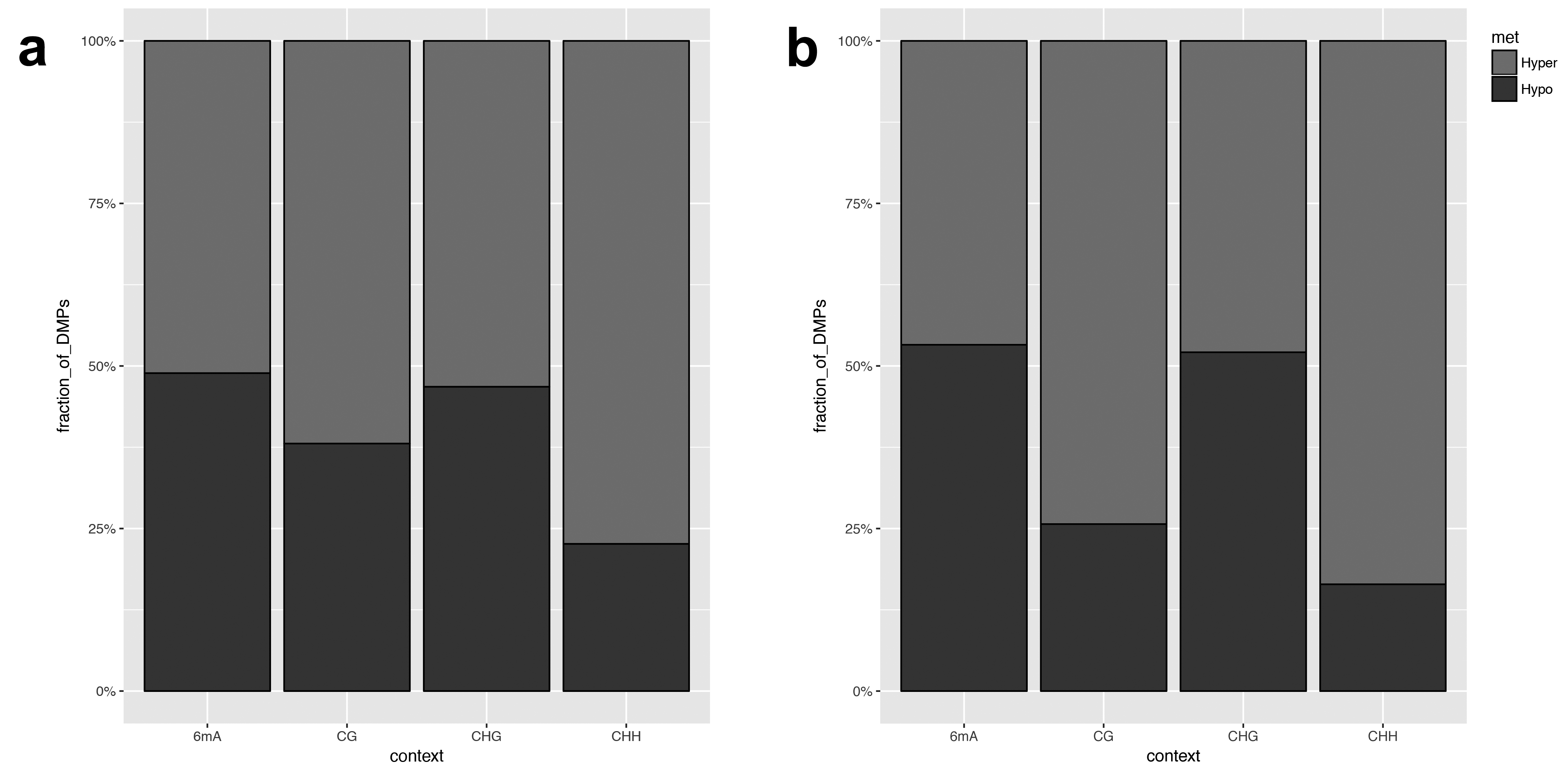

### Supplementary Figure 4

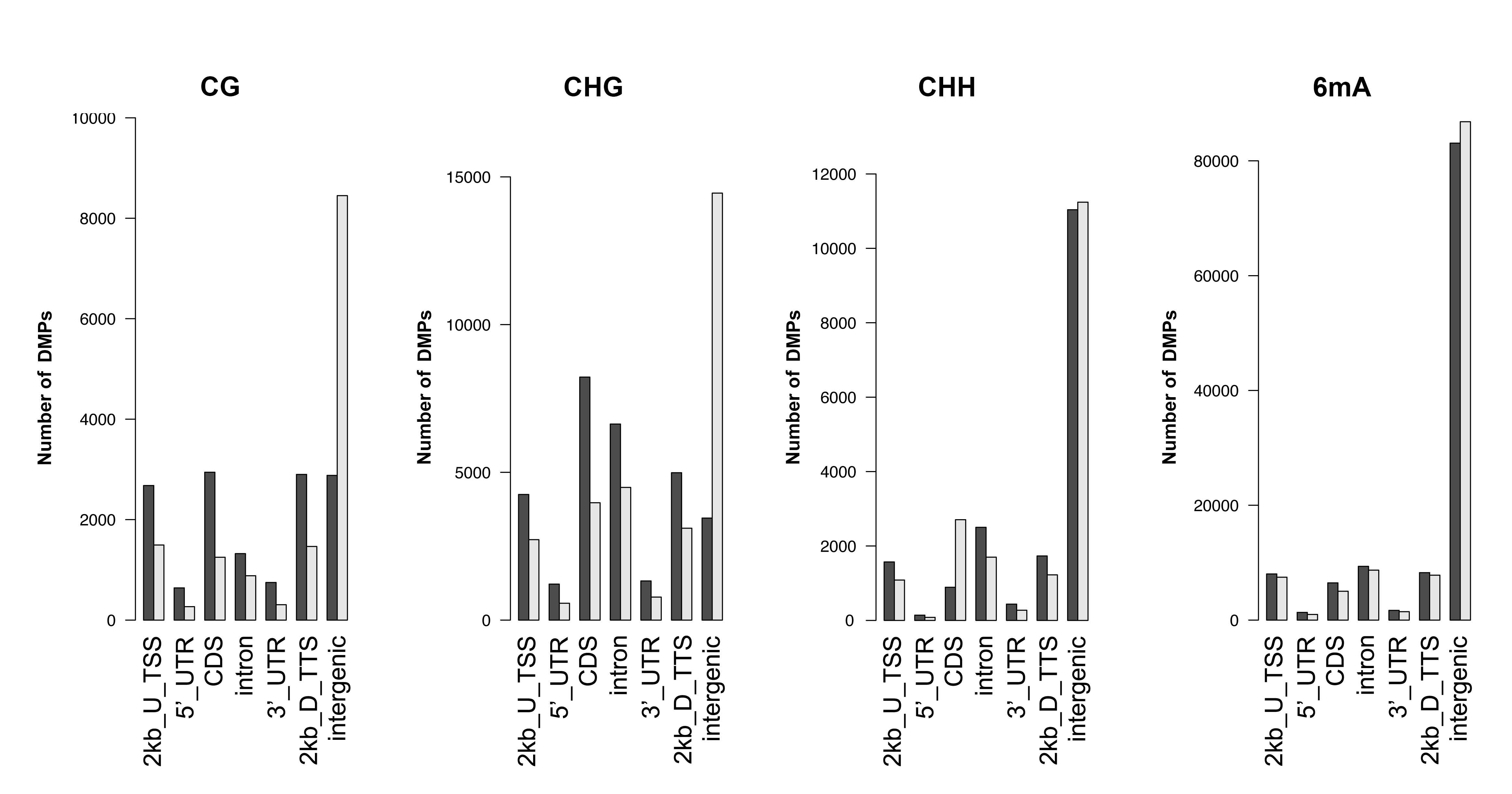

### Supplementary Figure 5

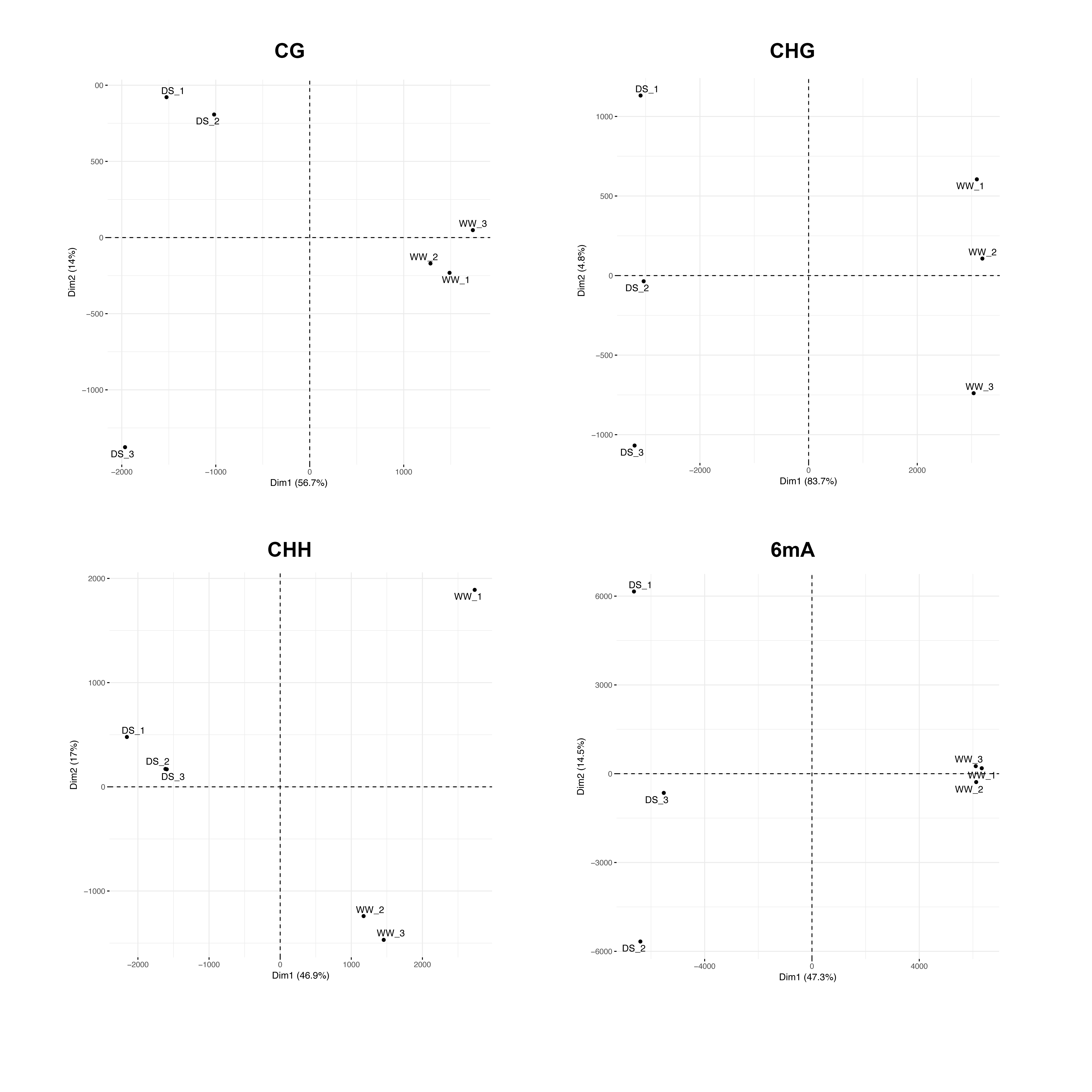

### Supplementary Figure 6

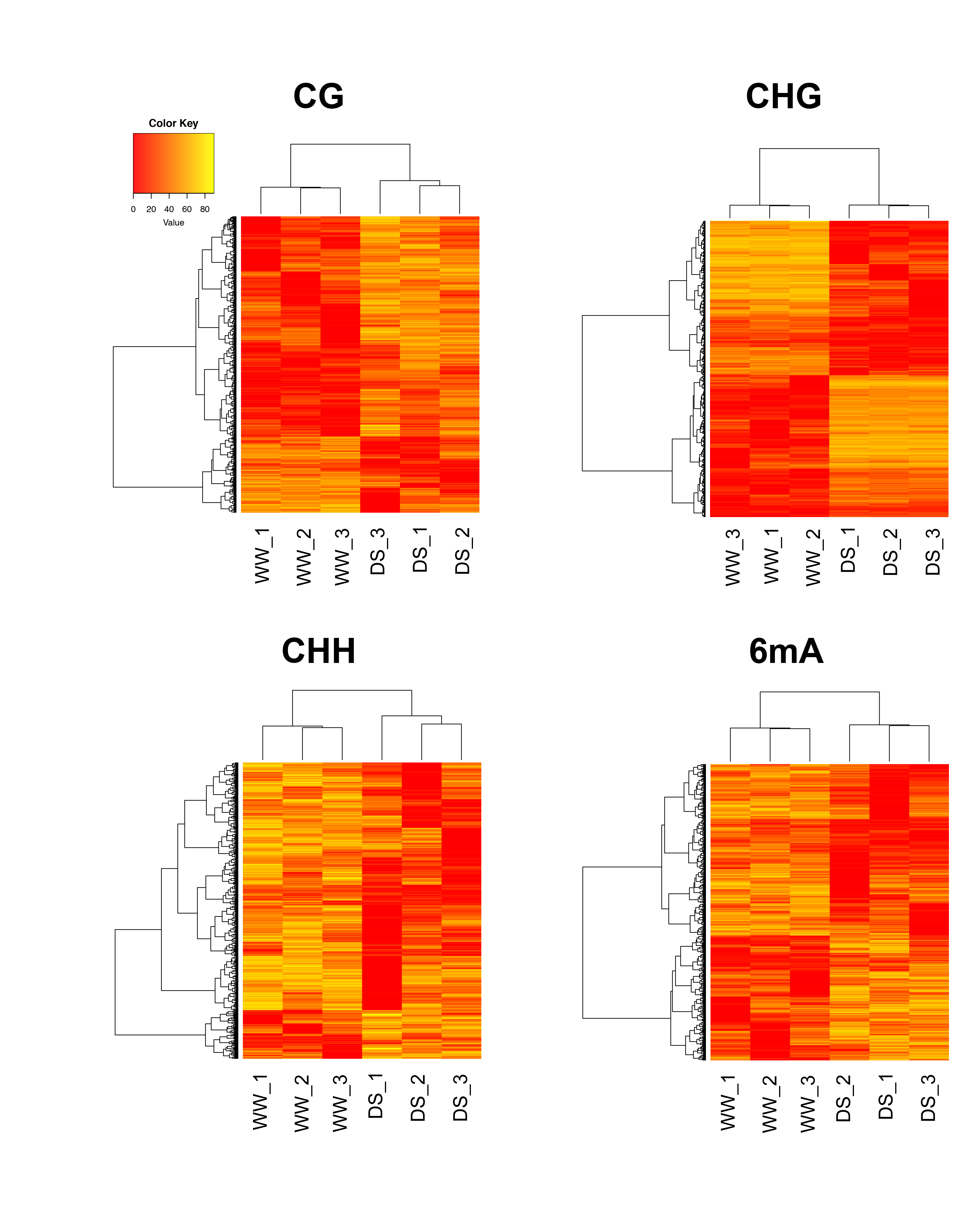
